# A RAF-like kinase mediates a deeply conserved, ultra-rapid auxin response

**DOI:** 10.1101/2022.11.25.517951

**Authors:** Andre Kuhn, Mark Roosjen, Sumanth Mutte, Shiv Mani Dubey, Polet Carrillo Carrasco, Aline Monzer, Takayuki Kohchi, Ryuichi Nishihama, Matyáš Fendrych, Jiří Friml, Joris Sprakel, Dolf Weijers

**Affiliations:** Laboratory of Biochemistry, Wageningen University, Stippeneng 4, Wageningen, the Netherlands; Department of Experimental Plant Biology, Charles University, Prague; Institute of Science and Technology Austria, Klosterneuburg, Austria; Graduate School of Biostudies, Kyoto University, Kyoto, Japan; Department of Applied Biological Science, Faculty of Science and Technology Tokyo University of Science, Noda, Chiba, Japan; Laboratory of Physical Chemistry and Soft Matter, Wageningen University, Wageningen, the Netherlands

**Keywords:** Auxin, protein phosphorylation, RAF kinase, plant evolution

## Abstract

The plant signaling molecule auxin triggers both fast and slow cellular responses across the plant kingdom, including both land plants and algae. A nuclear response pathway mediates auxin-dependent gene expression, and controls a range of growth and developmental processes in land plants. It is unknown what mechanisms underlie both the physiological responses occurring within seconds, and the responses in algae, that lack the nuclear auxin response pathway. We discovered an ultra-fast proteome-wide phosphorylation response to auxin across 5 land plant and algal species, converging on a core group of shared target proteins. We find conserved rapid physiological responses to auxin in the same species and identified a RAF-like protein kinase as a central mediator of auxin-triggered phosphorylation across species. Genetic analysis allowed to connect this kinase to both auxin-triggered protein phosphorylation and a rapid cellular response, thus identifying an ancient mechanism for fast auxin responses in the green lineage.

## INTRODUCTION

The plant signaling molecule is key to numerous growth and developmental processes in plants^1^. Iconic auxin-dependent processes are the tropic growth responses to light and gravity^2–5^, differentiation of vascular strands and the control of fruit development^6–9^. The dominant naturally occurring auxin is indole 3-acetic acid (IAA), a chemically simple Tryptophan derivative that land plants can synthesize in a two-step pathway, but that is widely found across both prokaryotic and eukaryotic species^10^. While initial discoveries with auxin were made in flowering plants, both the occurrence of IAA and physiological and developmental responses to the molecule have been reported well beyond this group. All land plants studied^11^, and a range of algae^12–14^ show responses to externally applied auxin, which suggests a very deep origin of the capacity to respond to auxin. The cellular responses to auxin come in essentially two flavors: fast and slow. The fast responses include changes in membrane polarization^15–17^, cytoplasmic streaming^18,19^, Calcium and proton fluxes^20–24^ and remodeling of the cytoskeleton^12,25^ and trafficking^26^. Slower responses include cellular growth, division and differentiation^27–30^.

Following an era of biochemical investigation that led to the identification of a set of auxin-binding proteins^31^, genetic approaches have been incredibly successful in defining a comprehensive response system. Using the ability of auxin to inhibit root growth in the flowering plant *Arabidopsis thaliana* as a model, a set of components was identified that mediates auxin’s activity in regulating gene expression – the nuclear auxin pathway (NAP)^32– 36^. This system revolves around the auxin-triggered proteolysis of a family of transcriptional repressor proteins, thus liberating DNA-bound transcription factors and allowing gene regulation^37^. Through this pathway, auxin controls the expression of hundreds-thousands of genes, and mutations in its components interfere with most, if not all developmental auxin functions, culminating in embryo lethality in the most affected mutants^38–40^.

As increasing numbers of plant genomes have become available, it became possible to reconstruct the occurrence and evolutionary history of the auxin response system. From such analysis, it appeared that the same auxin response system acts to control gene expression and development across land plants^11,41^. However, it is also clear that the closest sister group to land plants – the streptophyte algae – do not carry the NAP, in cases even lacking all its components^11^. Thus, a major unanswered question is how algae can respond to auxin in the absence of the known auxin response system. In addition, the fastest gene expression responses to auxin have been recorded in 5-10 minutes^42,43^, but several of the fast responses^18,19,23,44,45^ occur within seconds, or at least well within the time needed for gene expression and protein synthesis. Thus, it is likely that the currently known auxin response system represents the “slow” branch, and that a separate, currently unknown system must exist to mediate fast responses. The existence of fast auxin responses in land plants and their algal sisters would predict such a system to be shared between these clades.

Building on the rich literature in animal signaling, we explored the hypothesis that regulated protein phosphorylation may represent a mechanism mediating fast auxin responses. In the accompanying article (Roosjen, Kuhn et al., accompanying manuscript) we demonstrate that auxin can trigger changes in protein phosphorylation well within 30 seconds, and that more than 2000 proteins are targeted by auxin-triggered phosphorylation within 10 minutes in Arabidopsis roots. Auxin-triggered phosphorylation targets numerous pathways, including those leading to changes in membrane polarity. Here, we asked if this novel auxin response may represent the elusive, deeply conserved mechanism underlying rapid cellular responses. We indeed find that auxin triggers rapid changes in protein phosphorylation in 5 different land plant and algal species, including a core set of conserved targets. We show that auxin has deeply conserved activity in accelerating cytoplasmic streaming and membrane polarity. Lastly, we identify a key RAF-like kinase that mediates auxin-triggered protein phosphorylation and control of fast cellular responses across species. This work thus identifies an ancient system for rapid responses to the auxin signaling molecule.

## RESULTS

### Identification of a deeply conserved, rapid, phosphorylation-based auxin response

To examine whether the rapid phosphorylation-based auxin response that we have identified in *Arabidopsis thaliana* roots (Henceforth: Arabidopsis; Roosjen, Kuhn et al, accompanying manuscript) is conserved beyond this species, we selected a set of phylogenetically distant species ranging from green algae to bryophytes for phosphoproteomic analysis. These included the streptophyte algae *Klebsormidium nitens* (Klebsormidium) and *Penium margaritaceum* (Penium) and the bryophytes *Marchantia polymorpha* (Marchantia) and *Physcomitrium patens* (Physcomitrium) in addition to the angiosperm Arabidopsis. This selection encompasses both early-diverging streptophyte algae (Klebsormidium) and a close sister to land plants (Penium; Zygnematophyceae), and covers two clades within the bryophytes: liverworts (Marchantia) and mosses (Physcomitrium). Notably, while sporophytic (root) tissue was sampled for Arabidopsis, gametophyte tissue was sampled for all other species. Thus, the suite of species not only spans phylogeny, but also haploid and diploid generations. All species were treated with the same concentration (100 nM) of the naturally occurring auxin Indole 3-Acetic Acid (IAA, auxin), followed by phosphopeptide enrichment after two minutes using the same experimental, mass spectrometry and analysis workflow that we describe in Roosjen, Kuhn et al. (accompanying manuscript). Strikingly, we find that two minutes of auxin treatment leads to large shifts in the phospho-proteome in all species tested (Figure 1A). The number of differential phosphosites was comparable across species (FDR≥1.301: n=1048 in Arabidopsis; n=670 in Physcomitrium; n=741 in Marchantia; n=719 in Penium; n=1231 in Klebsormidium). In all species except Klebsormidium, hyperphosphorylation upon auxin treatment represented the majority of differential phosphosites (64% in Arabidopsis, 76% in Physcomitrium, 73% in Marchantia and 60% in Penium), while hyper-and hypophosphorylation were more equal in Klebsormidium (47% hyperphosphorylation) (Figure 1A). Thus, rapid, global changes in phospho-proteomes are triggered by auxin at comparable scale in all species tested.

**Figure 1.**
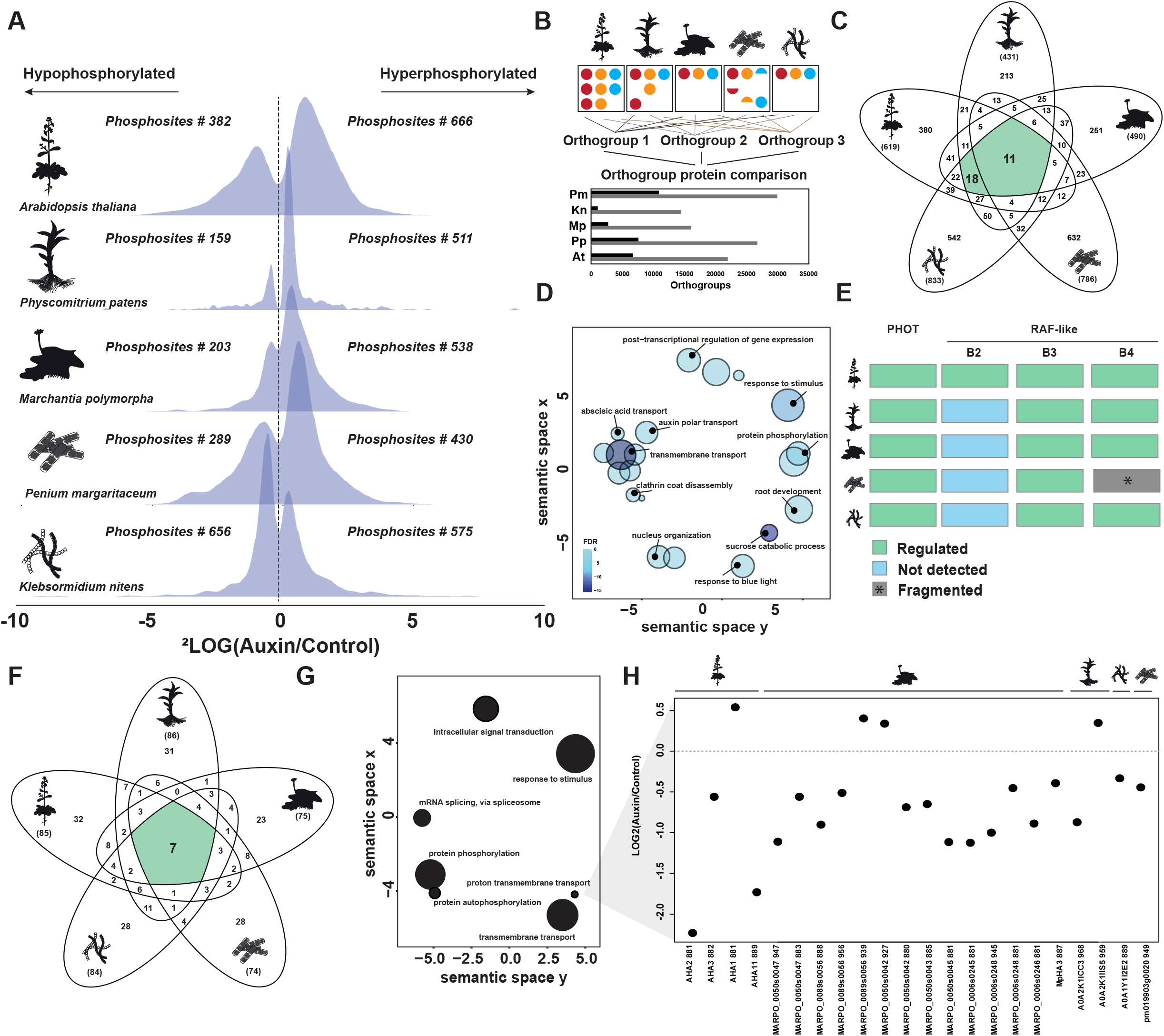
Comparative phosphoproteomics identifies a rapid and conserved auxin response. (A) Distribution histograms of significant differential phosphosites (FDR≤0.05) comparing 2 minutes of 100 nM IAA (Auxin) treatment with mock treatment across 5 species. Numbers of hyper- or hypo-phosphorylated sites are indicated. (B) Strategy for orthogroup based on protein sequence across the 5 species used here (top). The lower panel shows the number proteins residing in shared (black) and unique (grey) orthogroups in each species. (C) Venn diagram depicting the orthogroups found as differentially phosphorylated upon auxin treatment in all 5 species. (D) Reduced GO analysis (Revigo) of the 29 shared orthogroups (marked green in panel C). Circle sizes correspond to gene count within orthogroups. (E) Heatmap depicting measured significantly differnential phosphosites (FDR≤0.05) of two kinase families, PHOT and RAF-like kinases. (F) Venn diagram depicting the number of shared GO terms across all species tested, based on closest Arabidopsis homolog of each differential protein (FDR≤0.05). (G) Reduced GO analysis (Revigo) of the 7 shared orthogroups (marked green in panel F). (H) Differential phosphorylation of plasma membrane H^+^-ATPases across all species tested.

We next asked if the cellular functions and proteins that are targeted by auxin-triggered phosphorylation changes are conserved among the species tested. Estimated divergence times of the species used here from common ancestors is around 850 Mya for algae and land plants, and 500 Mya among the land plants^46^. Given these enormous evolutionary distances, there is substantial sequence divergence within protein families, and large differences in gene family numbers^47^. This makes establishing direct orthology relationships very challenging. Therefore, before comparison of differential phosphoproteins at protein/family level, we first constructed a set of orthogroups that represent the set of genes that originated from a single gene in the last common ancestor of all the species under consideration. We then consider members of the same orthogroup to represent a conserved ancestral function. Among the species tested, Penium has a remarkably large number of orthogroups with multiple members within Penium (Figure 1B), which is a reflection of the high degree of fragmentation of the genome assembly^48^.

Comparing the phosphosites in all species, we found an overlap of 11 orthogroups across all organisms (Figure 1C). Given the previous consideration, we also consider orthogroups not represented in Penium to be relevant. When excluding Penium from the analysis we found 29 orthogroups to be shared (Figure 1C). Gene Ontology (GO) analysis on the conserved orthogroups showed that a broad range of cellular functions is subject to auxin regulation (Figure 1D). These include processes at the plasma membrane or endomembranes, such as transmembrane transport and clathrin coat disassembly, but also nuclear organization and posttranslational regulation of gene expression. Furthermore, GO analysis identified responses to external stimuli and hormones, including response to blue light, abscisic acid transport and polar auxin transport. As expected from a phospho-proteomic analysis, protein phosphorylation was another highly enriched GO-term. In line with that, we find RAF-like kinases and the blue-light receptor PHOT1 as a conserved target of auxin-triggered phosphorylation (Figure 1E).

Limiting GO analysis to only the 29 conserved orthogroups is very stringent, as it is strongly constrained by sequence similarity, which may be limited across such long evolutionary timescales. We therefore also performed GO analysis on the full set of differentially phosphorylated phosphosites (FDR≤0,05) in each species separately, and compared the enriched GO-terms. This comparison found 7 GO-terms enriched in all species tested (Figure 1F), suggesting that these represent core target processes of rapid auxin response. Beyond the previously identified GO terms (Figure 1D), “transmembrane transport” and “proton transmembrane transport” were highly enriched (Figure 1G). Further analysis showed that in all species tested, H^+^-ATPase proton pumps were differentially phospho-regulated upon auxin treatment (Figure 1H). Clearly, there were also many GO-terms that were uniquely enriched in one or a few species (Figure 1F), suggesting that rapid auxin-triggered phosphorylation not only has a conserved component, but also a species/lineage-specific component. In conclusion, auxin triggers a conserved set of rapid phosphorylation changes across land plants and algae, converging on shared cellular processes.

### Auxin triggers fast cellular and physiological responses across the plant lineage

The identification of a deeply conserved auxin response that targets a common set of proteins and functions, suggests that there are cellular processes that auxin can regulate across the green kingdom. Among the fast auxin responses that have previously been recorded (see introduction), two stand out as being potential candidates for being shared outside of land plants. We explored whether auxin can trigger changes in membrane polarity and cytoplasmic streaming across species.

Membrane potential reflects the difference between cytoplasmic and apoplastic electrical potentials (Figure 2A). Auxin has a profound effect on membrane potential by triggering instantaneous depolarization of plasma membranes. This depolarization is then followed by a hyperpolarization of the membrane^17,24^. Both membrane depolarization and hyperpolarization depend on auxin’s ability to regulate ion fluxes across the plasma membrane, prominently involving H^+^-ATPase proton pumps^17,23,49,50^. To test whether this response is conserved in the plant lineage, we monitored membrane potential after 5 min of treatment with 100 nM auxin in Arabidopsis roots, Marchantia gemmae and Klebsormidium filaments using the membrane potential fluorescent probe DISBAC_2_(3) ^17,51^. Increase in DISBAC_2_(3) fluorescence reports membrane depolarization^17,51^. We observed a significant increase of fluorescence ratio upon auxin-treatment in all three species (Figure 2A). Moreover, the increase was quantitatively very similar between species. This indicates that rapid auxin-triggered plasma membrane depolarization is a deeply conserved rapid auxin response.

**Figure 2.**
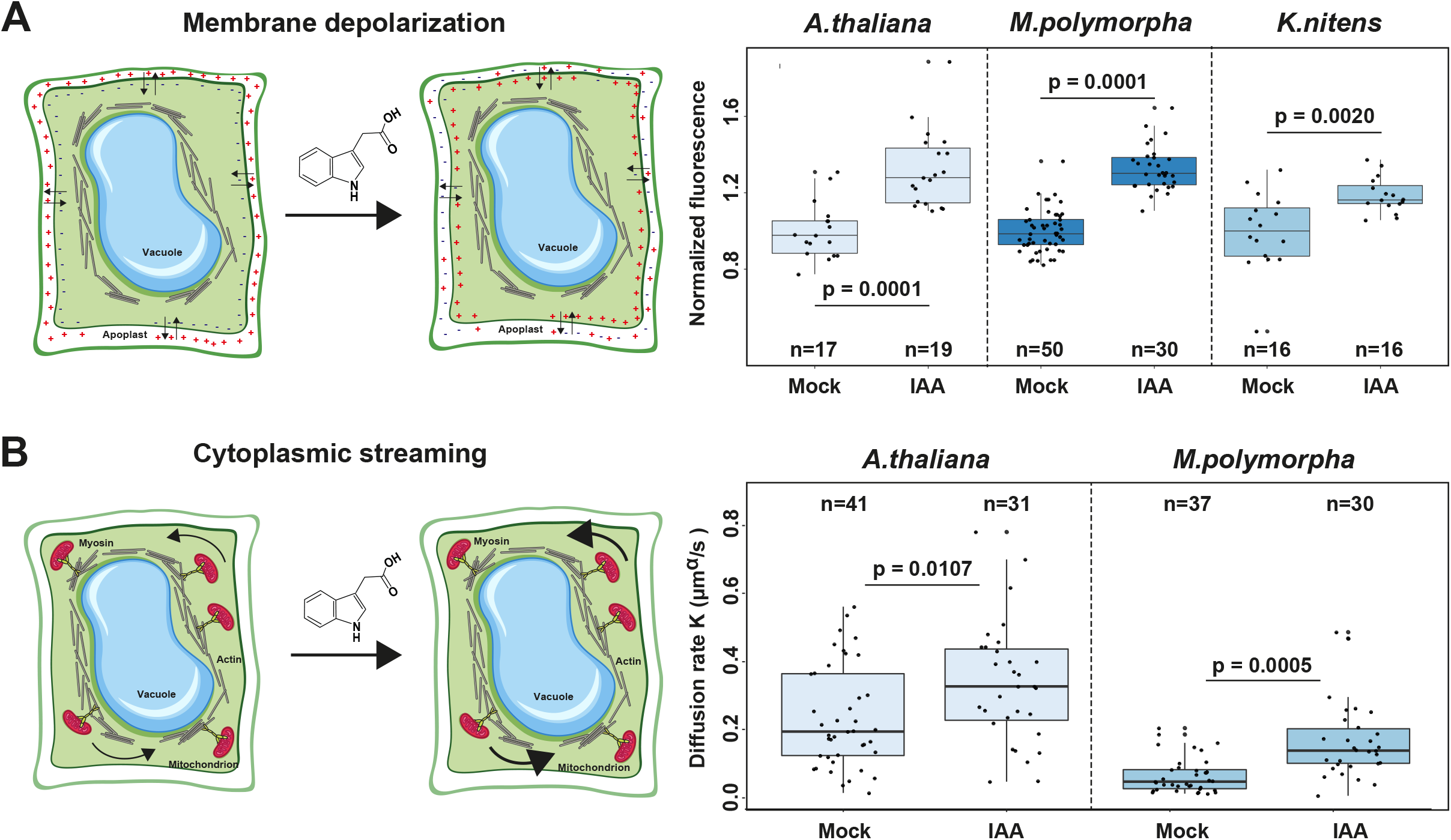
Auxin triggers fast cellular and physiological responses across the plant kingdom. (A) Scheme depicting membrane polarity and depolarization measured using DISBAC2(3) fluorescence (left) and normalized fluorescence in control (mock) and IAA-treated Arabidopsis root cells, Marchantia thallus cells and Klebsormidium cells. (B) Scheme depicting cytoplasmic streaming (left) and diffusion rate K (μm^α^/s) in control (mock) and IAA-treated Arabidopsis root cells and Marchantia thallus. Boxplots are shown along individual measurements, number of observations (n) is indicated, and significance (Student’s t-test) is shown.

Cytoplasmic streaming describes the movement of organelles along the actin cytoskeleton and is thought to have essential function in transport of nutrient and proteins within the cell ^52^. In plants, cytoplasmic streaming is thought to be primarily driven by plant-specific Myosin XI cytoskeletal motor proteins^52^. We found that in Arabidopsis, Myosin XI-K and the MadB Myosin-binding proteins are targets of rapid auxin-dependent physophorylation^53^, and that auxin promotes cytoplasmic streaming in root epidermal cells^18^. We examined the physiological effect of 100 nM auxin on cytoplasmic streaming by monitoring the movement of fluorescently labeled mitochondria in epidermis cells within the root elongation zone in Arabidopsis and in Marchantia rhizoid cells (Figure 2B). After particle tracking, we determined the active diffusion rate (K) and diffusive exponent (α) by fitting mean-square displacements, ensemble-averaged per cell, to the anomalous diffusion model, in both auxin treated and untreated samples. We detected consistent streaming within both species, but found absolute rates to differ among species (Figure 2B). Pretreatment of Arabidopsis roots with the actin depolymerizing drug Latrunculin B reduced cytoplasmic streaming in both species (Supplementary Figure 1B), thus implicating the actin cytoskeleton. Importantly, auxin treatment increased the diffusion rate in all species tested (Figure 2B). Hence, like membrane depolarization, acceleration of cytoplasmic streaming is a deeply conserved cellular response to auxin.

### Identification of RAF-like kinases as conserved components in auxin response

The finding that there are conserved phosphorylation responses to auxin in algae and land plants, along with conserved cellular responses, suggests the existence of a shared mechanism for auxin perception and signal transduction. Given the prominent phosphorylation changes across species and the temporal dynamics of the response in Arabidopsis (Roosjen, Kuhn et al, accompanying manuscript), we anticipate a key role for auxin-activated protein kinases. To identify such kinases, we first analyzed phosphorylation motifs enriched among the conserved phospho-targets. We found that hyperphosporylation was associated with the presence of a proline-directed SP motif (Figure 3A). These are typically targeted by MAP kinases^54,55^. Indeed, when inferring kinase-target networks in Arabidopsis from temporal phosphorylation profiles and activation loop phosphorylation predictions, we identified an auxin-activated RAF-like Kinase as a potential hub, central to the phosphorylation network (Roosjen, Kuhn et al., accompanying manuscript). Strikingly, orthologues of this same Rapidly Accelerated Fibrosarcoma (RAF)-like kinase were hyperphosphorylated upon auxin treatment in all other species tested (Figure 1E and 3B), except in Penium, where genome assembly fragmentation likely precluded its identification. In addition to the RAF-like kinase, we also identified PHOT1 as a conserved target of auxin-triggered hyperphosphorylation (Figure 1E). However, given the multiple lines of evidence suggesting a role for the RAF-like Kinases in auxin-triggered phosphorylation, we here focus on this protein.

**Figure 3.**
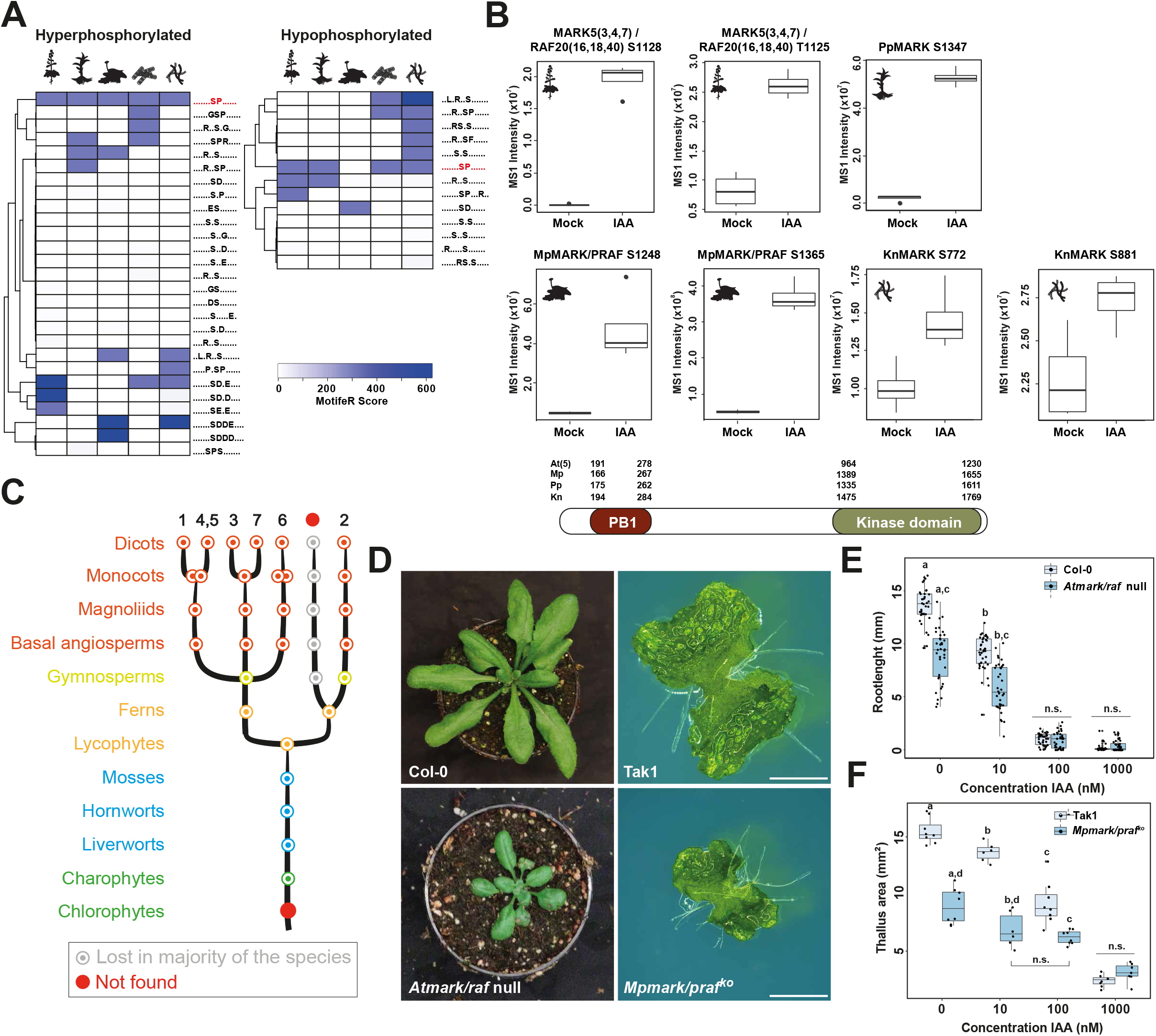
Identification of MARK/RAF-like kinases. **(A)** Clustering of phosphomotif enrichment scores (using motifeR) of significantly differential (FDR≤0.05) phosphosites in all tested species. **(B)** Raw MS1 intensities of RAF-like kinase orthologues in mock- and IAA-treated samples. Phosphorylated residues are indicated. Lower: domain topology of B4 RAF-like kinases indicating positions of PB1 and kinase domains (residue numbers) in Arabidopsis MARK5/Raf24 (At), Marchantia MARK/PRAF (Mp), Physcomitrium MARK (Pp) and Klebsormidium MARK (Kn). **(C)** Inferred phylogeny of the B4 RAF-like kinase. Arabidopsis numbering is indicated on the top. Every node represents an inferred ancestral gene copy at each divergence event. The complete tree can be found at interactive Tree of Life (iTOL): https://itol.embl.de/shared/dolfweijers. **(D)** Phenotype of Arabidopsis (left) Col-0 wild-type and *mark/raf* null mutant rosettes and Marchantia (right) Tak-1 wild-type and *mark/praf* mutants thallus **(E**,**F)** Length of Col-0 wild-type and *mark/raf* mutant Arabidopsis roots (**E**) and area of Tak-1 wild-type and *mark/praf* mutant Marchantia thallus (**F**) on increasing concentrations of IAA. Distributions at each concentration were tested for significant differences using ANOVA.

RAF-like kinases are serine/threonine kinases that belong to the mitogen activated protein kinase kinase kinases (MAPKKKs) family. They are classified into four B clades and seven C clades according to their homology with the widespread eukaryotic RAF protein kinases^56^. Arabidopsis B2, B3 and B4 clade RAF-like kinases have been implicated in various physiological responses, including responses to hypoxia, osmotic stress and drought^57,58^. The Marchantia B4 RAF-like kinase (PRAF) was implicated in the regulation of carbon fixation^59^. While we found RAF-like kinases of the B2, B3 and B4 clade to be hyperphosphorylated after auxin treatment in Arabidopsis, it seems that only hyperphosphorylation of RAF-like kinases of the B3 and B4 clade upon auxin treatment is conserved (Figure 1E). The B4 clade is represented by 7 paralogs in Arabidopsis, 2 in Physcomitrium and single copies in Klebsormidium and Marchantia^60^ (Figure 3C). Most of these are hyperphosphorylated in response to auxin treatment (Figure 3B), firmly connecting this family to auxin response. We refer to these proteins as MAP AUXIN RESPONSIVE KINASE/RAFs (MARK/RAFs).

Given that no role for these proteins in auxin response has been reported, we initially explored requirements for MARK/RAF kinases in auxin-associated growth and development, as well as in response to externally applied auxin. To this end, we analyzed previously established mutants: two septuple mutants of the entire Arabidopsis B4 clade either conferring a null (*mark/raf*^null^; also referred to as OK^130^-*null* ^58^) or a weak allele combination (*mark/raf*^weak^; also referred to as OK^130^-*weak*^58^), and a null mutant in the single Marchantia ortholog (Mp*mark/praf*^KO^, also referred to as Mp*praf*^KO 59^). We found that in both species, loss of MARK activity caused growth and developmental phenotypes (Figure 3D). While in Arabidopsis we found a range of defects in root growth, plant height and rosette area and germination (Figure 3D; Supplementary Figure 2), in Marchantia, these manifested as smaller thallus size and reduced gemmae cup number confirming previously published results^59^ (Figure 3D; Supplementary Figure 2). Essentially all these processes are known to involve auxin action^30,61,62^. We therefore tested sensitivity of the Arabidopsis and Marchantia *mark/praf* mutants to auxin. In Arabidopsis, *mark* mutant roots were slightly less sensitive to growth inhibition by auxin (Figure 3E). Likewise, Marchantia *mark/praf* mutant thallus, although already under control conditions reduced in size, was also less sensitive to auxin-induced growth inhibition (Figure 3F). Thus, in both species, MARK/RAF kinases act in growth and development, and play a role in auxin response.

### MARK kinases mediate fast auxin phospho-response

Auxin-associated growth and development, as well as Arabidopsis root and Marchantia thallus growth responses to externally applied auxin is typically associated with changes in auxin-dependent gene expression through the NAP^37,41^. given the auxin-related phenotypes in *mark* mutants, we asked if these are affected in transcriptional responses. We therefore performed RNA-Seq in Arabidopsis (roots) and Marchantia (thallus) wildtype and *mark/raf* mutants that were either treated with 1μM IAA or control medium for one hour. This concentration of IAA should allow to detect even subtle changes in transcription in mutants. In both species, transcriptomes under untreated conditions look very distinct between mutant and wildtype (Figure 4A,B), suggesting massive effects of loss of MARK/RAF function on the “baseline” transcriptome in the absence of externally applied auxin. However, comparing auxin-treated and untreated samples in both species showed substantial auxin-induced changes in transcriptomes in both wildtypes and in *mark/raf* mutants (Figure 4A,B). Qualitatively, mutants in both species still showed a typical gene expression response to auxin. Indeed, detailed analysis of individual auxin-regulated genes (Figure 4A,B) showed that mutants did not have an obvious defect in auxin-induced transcription. This suggests that MARK/RAF proteins do not have a major role in transcriptional auxin responses.

**Figure 4.**
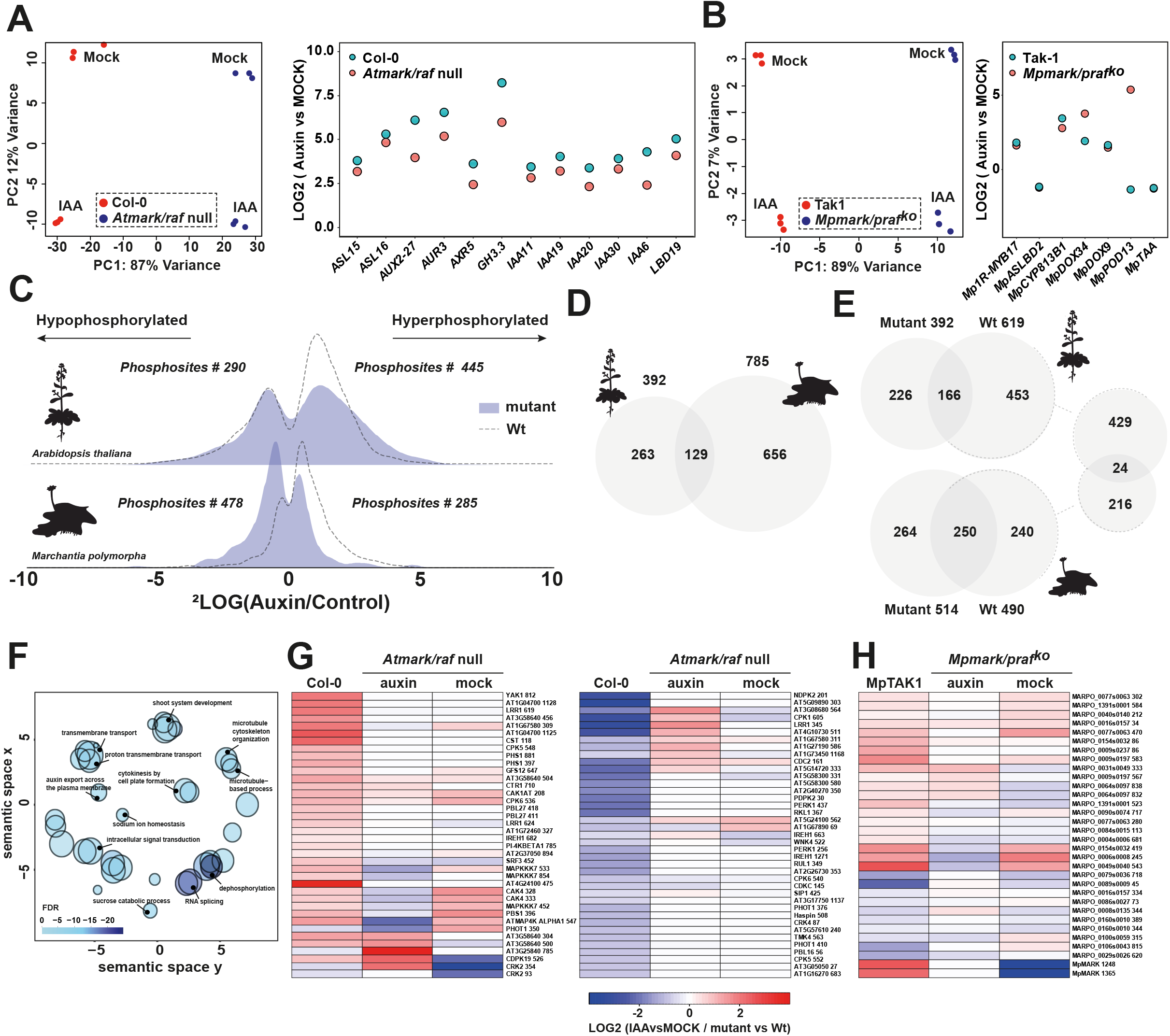
MARK mediates auxin posphoresponse across land plant species. **(A**,**B)** PCA plots (left) and expression analysis of individual, auxin-regulated genes (right) from RNA-seq analysis on (Col-0; Tak-1) wildtype and *mark* mutants in Arabidopsis roots and Marchantia gemmae treated with 1μM IAA for 1 hour. **(C)** Distribution histograms of significant differential phosphosites (FDR≤0.05) comparing 2 minutes of 100 nM IAA (Auxin) treatment with mock treatment in wild-type (dashed lines) and mark mutant (solid area) Arabidopsis roots (top) and Marchantia gemmae (bottom). Number of phosphosites is indicated. **(D)** Venn diagrams indicating orthogroup overlap of significantly differential phosphosites (FDR≤0.05) in *mark* mutants in Arabidopsis and Marchantia compared to respective wild-types under mock condition. **(E)** Venn diagrams indicating orthogroup overlap of significantly differential phosphosites (FDR≤0.05) in *mark* mutants and wild-types in Arabidopsis and Marchantia under IAA-treated condition. **(F)**. Gene ontology analysis on the overlapping and conserved auxin- and MARK-dependent proteins. (**G**,**H**) Heatmap showing differential phosphorylation in Arabidopsis (**G**) and Marchantia (**H**) *mark* mutants of all kinases that are auxin-regulated in wild-type.

Given the rapid activation of MARK/RAF kinases by auxin (Figure 3B), it is conceivable that these kinases act in auxin response through their role in mediating rapid phosphorylation responses. We tested this hypothesis by subjecting *mark/raf* mutants in both Arabidopsis and Marchantia to phosphoproteomic profiling after two minutes of treatment with 100 nM IAA or control media. In both species, we found that the number of significant differential hyperphosphorylated phosphosites after auxin treatment was reduced (666 in Arabidopsis WT; 445 in At*mark/raf* mutant; 538 in Marchantia WT; 285 in Mp*mark/praf*; Figure 4C). When comparing the number of phosphosites in wild-types and mutants, we found that 73% of the differential phosphosites in wild-type was lost in the Arabidopsis *mark/raf* mutant, while 51% was lost in the Marchantia *mark/praf* mutant (Figure 4C). We compared phosphoproteomes in non-treated mutants with wild-type controls in both species to identify functions that are deregulated in *mark* mutants. In Arabidopsis *mark/raf*, 392 orthogroups were different between mutant and wildtype, while in Marchantia *mark/praf*, 785 orthogroups were differentially phosphorylated (Figure 4D). Many orthogroups that were not significantly affected by auxin in wild-type became differentially phosphorylated upon auxin treatment in the mutants (Figure 4E). This suggests that the mutants in both species not only lack a substantial part of auxin-triggered phosphorylation, but also have a response system that is differently wired in non-treated conditions. This is consistent with the large transcriptional changes, and with the strong phenotypes in the mutants. When comparing targets of MARK-dependent, auxin-triggered phosphorylation changes in the two species, we found a small overlap (24 orthogroups; Figure 4E). Given the evolutionary distance between Marchantia and Arabidopsis, this is remarkable since it suggests that there is indeed a set of evolutionary conserved fast auxin response under control of a conserved mechanism. These shared, MARK/auxin-dependent targets included proteins associated with a diverse set of cellular processes (Figure 4F; Supplementary Figure 3A). This includes ion transport, membrane dynamics, and auxin export (e.g. PIN’s, ABCB’s, D6PK), but also featured nuclear processes such as splicing and cytoplasmic processes such as cell plate formation and cytoskeleton organization (e.g. SPIKE1, TOR1, NEK5). Lastly, this analysis also identified previously reported phospho-targets of B4-type RAF kinases (e.g. VCS, VCR, SE).

To explore to what extent the auxin-triggered phosphorylation network is affected in *mark* mutants, we compared the phosphorylation state of all kinases that were significantly hypo- or hyperphosphorylated upon auxin treatment in wild-type of both species with their phosphorylation state in *mark* mutants. Notably, in *mark* mutants, most of the auxin-triggered kinase phosphorylation was lost (Figure 4G,H). This suggests that MARK/RAFs directly or indirectly regulates the auxin-triggered phosphorylation of these kinases.

### Specificity and mechanism of MARK activation

Proteins in the Arabidopsis MARK/RAF family have been identified as being hyperphosphorylated upon osmotic treatment^58^ and to mediate response to hypoxia^57^, while Marchantia MARK/PRAF has a role in the response to altered photosynthesis^59^. This suggests that the same kinase is part of multiple response pathways and urges the questions of how specific the auxin-triggered phosphorylation changes are, and how MARK/RAF is activated in the context of auxin response. We initially compared the 2-minute auxin-triggered phosphorylation changes with the set of 973 phosphosites that are osmotic stress-responsive in Arabidopsis^58^. The overlap was very limited (37 phosphosites; Supplementary Figure 3B), and 13 of these overlapping phosphosites depend on MARK/RAF (Supplementary Figure 3B). We therefore conclude that the phosphoresponse that we identified here is specific and independent from osmotic stress responses.

We next explored mechanisms of MARK/RAF activation. In time-course phosphoproteome data (derived from Roosjen, Kuhn et al., accompanying manuscript), we found that multiple sites on all AtMARK/RAF proteins are modulated upon auxin treatment (Figure 5A), suggesting profound and rapid regulation. As part of our characterization of the auxin-triggered fast phosphoproteome in Arabidopsis, we found that the ABP1 auxin binding protein and the TMK1 receptor-like kinase as well as the intracellular AFB1 receptor contribute to effects of auxin on the phosphoproteome (Friml et al., 2022; Roosjen, Kuhn et al., accompanying manuscript). We found that phosphoproteome changes in *abp1* and *tmk1* mutants are highly correlated (Roosjen, Kuhn et al., accompanying manuscript), while effects on the same phosphosites in *afb1* mutants often are anticorrelated (Roosjen, Kuhn et al., accompanying manuscript). We compared the Arabidopsis *mark* mutant phosphoproteomes with those of wild type, *afb1, abp1* and *tmk1* mutants and found that phosphosites in *mark/raf* phosphoproteomes overlap less with those of *afb1*-3 and the auxin-treated wildtype phosphoproteome than with those of *tmk1* and *abp1* (Figure 5B,D). This is true for both *mark* phosphoproteomes in control and auxin-treated conditions. suggesting that the *mark* phosphoproteome under mock conditions is already strongly distorted. Given that MARK/RAFs do not have a clear ligand-binding domain, we were interested to see if MARK/RAFs phoshorylation depends on ABP1/TMK1 and/or AFB1. Therefore, we arrayed all phosphosites in AtMARK’s and compared their phosphorylation state in the mutant backgrounds (Figure 5C). In this analysis it is clear that MARK/RAF phosphorylation is strongly disturbed in each mutant, and that the *afb1* pattern more closely resembles that of wild-type, whereas *abp1* and *tmk1* more severely disturb MARK/RAF phosphorylation (Figure 5C). Interestingly, consistent with global patterns of the entire phosphoproteome (Roosjen, Kuhn et al, accompanying manuscript), for some MARK/RAF sites, phosphorylation is antagonistically distorted between *afb1* mutants and *abp1* and *tmk1* mutants.

**Figure 5.**
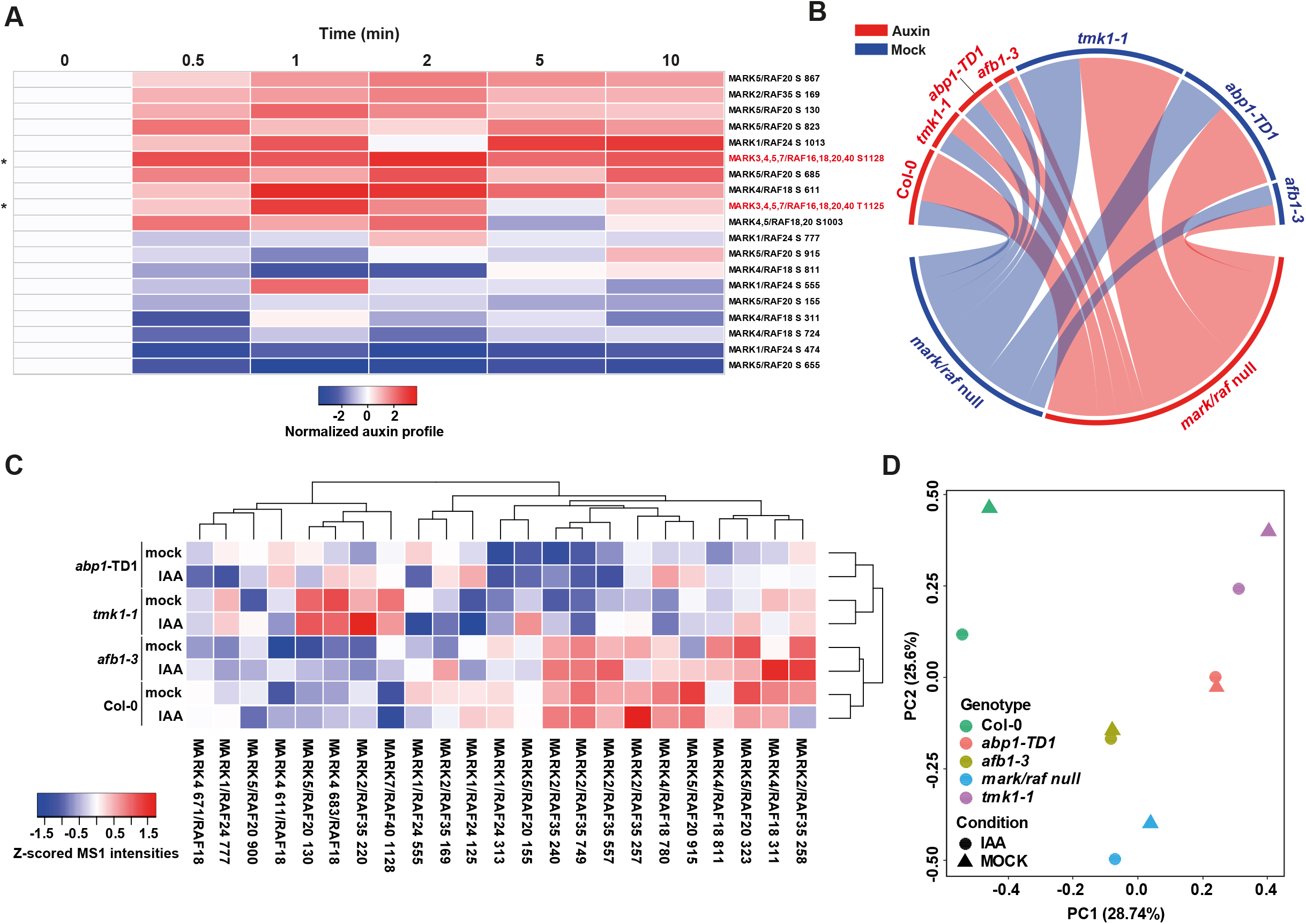
Requirements of MARK activation and activity. **(A)** Heatmap showing phosphorylation profiles, normalized to the t=0 timepoint, of Arabidopsis MARK/RAF kinases (data from Roosjen-Kuhn et.al., accompanying manuscript). Profiles marked with asterisk and red name are phosphosites located in the activation loop. **(B)** Chord plot depicting overlap between significant (FDR≤0.05) phosphosites in Arabidopsis mutants challenged with auxin (red) or without (blue). Overlap shows that the *mark/raf* mutant shares more commonly regulated phosphosites with *tmk1-1 and abp1-TD1* mutants than with the *afb1-3* mutant. **(C)** Z-scored MS1 intensities off all measured phosphosites of Arabidopsis MARK/RAF kinases in wild-type, *afb1-3, tmk1-1* and *abp1-TD1* mutants with or without IAA. **(D)** Principal component analysis of Z-scored MS1 intensities of all 1048 phosphosites that are auxin-regulated in wild-type in control- and auxin-treated wildtype, *tmk1-1, abp1-TD1 and afb1-3* mutants.

### MARK links rapid phospho-response to fast auxin responses

MARK/RAF proteins are a unique family of kinases that carry an N-terminal Phox-Bem1 domain (PB1)^63^ in addition to their C-terminal kinase domain (Figure 3B). PB1 domains can either mediate heterotypic or homotypic protein interaction with other PB1 domains, which can assemble into dimers or oligomers^64,65^. Apart from a single Arabidopsis paralog (HCR1)^57^, MARK/RAF protein localization has not been studied. We therefore generated translational fusions of Arabidopsis and Marchantia MARK/RAF proteins to fluorescent proteins and determined their localization. In both species, MARK/RAF proteins localized to punctate structures (Figure 6A, B) resembling the “punctae” observed for other PB1-containing proteins^66–68^. In both Arabidopsis roots (Figure 6A) and Marchantia gemmae (Figure 6B), these structures were associated both with the plasma membrane and in the cytoplasm. Thus, MARK/RAF protein locates to sites where fast auxin responses occur.

**Figure 6.**
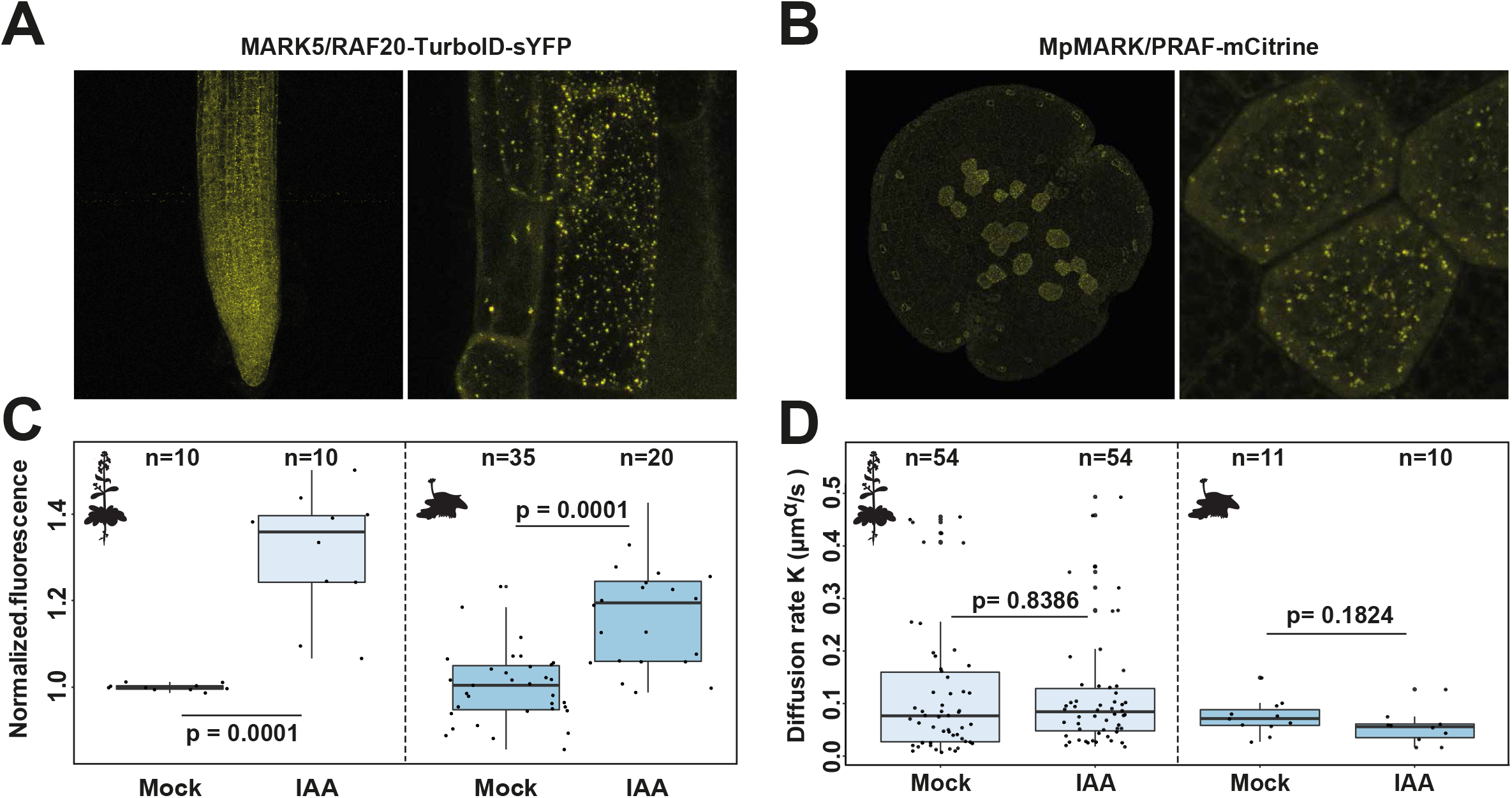
MARK links rapid phospho-response to fast auxin responses. (**A**) Fluorescence of Arabidopsis MARK5/RAF20-TurboID-sYFP, driven from its endogenous promoter, in primary root tips. Right panel shows close-up of epidermal cells. **(B)** Fluorescence of Marchantia MARK/PRAF-Citrine driven from its endogenous promoter in gemma. Right panel shows close-up of rhizoid initial cells. **(C)** Analysis of membrane depolarization on Arabidopsis and Marchantia *mark* mutants in mock and IAA-treated root (Arabidopsis) and thallus (Marchantia) cells (compare to Figure 2 A for wild types). Displayed is the normalized DISBAC2(3) fluorescence (IAA/mock). **(D)** Cytoplasmic streaming in Arabidopsis and Marchantia *mark* mutants in mock and IAA-treated root (Arabidopsis) and thallus (Marchantia) cells (compare to Figure 2B for wild types). Displayed is the Diffusion rate K (μm^α^/s). Boxplots are shown along individual measurements, number of observations (n) is indicated, and significance (Student’s t-test) is shown.

Given the profound role of MARK/RAF in mediating fast auxin-triggered phosphorylation changes, we explored whether MARK/RAF might mediate the rapid effect of auxin on membrane potential and cytoplasmic streaming. Responses to auxin treatment in membrane depolarization were normal in *mark* mutants in both Arabidopsis and Marchantia (Figure 6C, Supplementary Figure 4). However, we did find that Arabidopsis *mark/raf* mutants showed an altered apoplastic root surface pH profile (Supplementary Figure 5), perhaps caused by altered developmental zonation. Nonetheless, MARK/RAF does not appear to mediate auxin-triggered membrane depolarization (Figure 6C, Supplementary Figure 4) and the root surface alkalization response (Supplementary Figure 5).

In contrast, already in untreated Arabidopsis *mark/raf* mutant root epidermal cells, cytoplasmic streaming is significantly reduced (Figure 6D; compare with Figure 2B). Interestingly, *mark/raf* mutants are essentially insensitive to the promoting effect of auxin in cytoplasmic streaming (Figure 6D). In Marchantia rhizoid cells, *mark/praf* mutants showed wild-type cytoplasmic streaming in untreated conditions, but like in Arabidopsis, mutant cells were insensitive to the promoting effect of auxin (Figure 6D; compare with Figure 2B). This suggests that MARK proteins have a conserved role in mediating auxin-promoted cytoplasmic streaming in Arabidopsis and Marchantia. Collectively, we conclude that MARK proteins link rapid phosphorylation changes to a fast cellular response to auxin.

## DISCUSSION

Over the past decades, there have been impressive advances in understanding how auxin is synthesized, transported and degraded, and how it controls plant growth and development by regulating gene expression^37^. There are however several major open questions. Firstly: there is a number of auxin responses that are too rapid to be mediated by gene regulation, for which there is no mechanism yet. Secondly, no known mechanism can account for responses to auxin in algae, that lack the well-known transcriptional auxin response system^11^. In the accompanying article (Roosjen, Kuhn et al., accompanying manuscript) we identify a fast, unknown and unsuspected branch of auxin activity based on rapid protein phosphorylation, in Arabidopsis roots. Here, we demonstrate that this pathway is conserved across the green lineage, extending beyond land plants into the streptophyte algae. We show that some fast cellular responses to auxin are also conserved across land plants and algae and identify a key protein kinase mediating both auxin-triggered phosphorylation and a rapid cellular response. This identifies rapid phosphorylation-dependent signaling as a mechanism that can account for both fast and deeply conserved auxin responses.

Although we compared phosphoproteomes in different tissue types, and in both sporophytic (for Arabidopsis) and gametophytic tissue (for all other species), we detected a core set of functions and orthologous protein groups that are shared between all. The most parsimonious explanation is that this core set represents a truly ancient auxin “regulome” that has been retained in all these species to serve core functions. This is not trivial, given the estimated divergence times of between 850-500 Mya. In addition to the core set, there are numerous lineage/clade/group/organism-specific targets. This suggests profound diversification and neo-functionalization of auxin-triggered phosphorylation pathways. We have compiled all phosphoproteomics data generated in this study in the AuxPhos webtool (https://weijerslab.shinyapps.io/AuxPhos; Roosjen, Kuhn et al., accompanying manuscript), to allow facile access.

Though mining both comparative phosphoproteomics, kinase-substrate inference from temporal series and motif analysis, we identified a family of B4 RAF-like kinases (MARK/RAFs). Exploring mutants in orthologous proteins in Arabidopsis and Marchantia, we could establish that MARK/RAF kinases are central to auxin-triggered phosphorylation, and to development and physiological and cellular auxin response. Curiously, transcriptional auxin responses are not impaired, which suggests that the rapid, phosphorylation-based pathway is mechanistically uncoupled from the nuclear auxin pathway. The mutants, even in the absence of auxin treatment, have dramatic phenotypes. It should however be kept in mind that members of the MARK/RAF family have been implicated in responses to other triggers (e.g. light, osmotic stress)^57–59^. Disruption of these responses likely also contribute to the strong phenotypes, and dedicated strategies will be required to deconvolute these roles.

Notably, regulation of most kinases that are differentially phosphorylated upon auxin treatment in wild-type Marchantia and Arabidopsis, is lost in *mark* mutants, suggesting that MARK/RAF may sit at the apex of a multi-tier phosphorylation network. Interestingly, RAF kinases, MARK/RAF orthologs in mammals, play an important role as master regulator of signaling cascades, for example in EGF signaling^65^. MARK phosphorylation upon auxin treatment occurs within 30 seconds in Arabidopsis (the earliest sampled timepoint; Roosjen, Kuhn et al., accompanying manuscript). Mammalian RAF kinases can be activated by phosphorylation within seconds to minutes after signal recognition^69,70^. Therefore, the kinetics of MARK/RAF activation is consistent with the phospho-activation of their orthologs in animal cells.

Mammalian RAF Kinases polymerize through their PB1 domain and localize in punctate structures in the cytoplasm to form so-called signalosomes^65,68^. Signalosomes are large supramolecular protein complexes that help increase avidity between signaling components. The formation of such signaling hubs and their association with receptors is crucial for signal transduction in some pathways^64^. Curiously, both Arabidopsis and Marchantia MARK/RAF proteins localize to punctate structures in the cytoplasm and at the plasma membrane. It will be interesting to see if these punctae are functional signalosomes, whether they form through PB1 domain oligomerization, and what other proteins they bring together.

Inspired by the finding that algae and land plants share a common set of auxin phosphotargets, we explored if there are also shared cellular and physiological responses. Indeed, cytoplasmic streaming is deeply conserved responses across land plants while membrane depolarization is deeply conserved across land plants and algae. Both are widespread cellular phenomena that are connected to for example cellular growth, nutrient distribution and acquisition^50,71,72^. It is not clear what function the auxin-regulation of these processes serves, but analysis of these responses in *mark/raf* mutants did help to show bifurcation of rapid auxin response mechanisms. While auxin-dependent acceleration of cytoplasmic streaming depended on MARK/RAF, membrane depolarization did not. Interestingly, *mark/raf* mutants already had lower streaming velocity in the absence of auxin treatment, suggesting the same pathway operates during normal development, likely mediating the response to endogenous auxin.

The differential roles of MARK/RAF in the regulation of cytoplasmic streaming and membrane polarity are conserved between Arabidopsis and Marchantia, suggesting a deep evolutionary split between these two functions. In Arabidopsis, auxin-triggered membrane depolarization was previously attributed to the cytoplasmic AFB1 auxin receptor^17^, but its mechanism of action is not yet clear. Interestingly, AFB1 is a late innovation specific to angiosperms, and auxin-triggered membrane depolarization is found in the alga Klebsormidium that does not carry any TIR1/AFB ortholog^11^. This raises the question how the auxin signal translates to membrane depolarization outside of the angiosperms. Apart of the MARK/RAF-family, we identified B3-clade RAF-like kinases and PHOT1 kinases as potential conserved hubs in the auxin phosphorylation. It will be interesting to see if these kinases play a role in regulating membrane depolarization. Interestingly, PHOT1 was previously shown to mediate a rapid blue light-triggered membrane depolarization in Arabidopsis^73^, making it a strong candidate.

A key question is how the auxin signal is perceived and transmitted onto MARK/RAF proteins, given that MARK/RAFs do not have a clear ligand-binding domain. The auxin response components ABP1, TMK1 and AFB1 all contribute to auxin-triggered phosphorylation changes in Arabidopsis (Roosjen, Kuhn et al., accompanying manuscript). MARK/RAF phosphorylation was disturbed in all three mutants, but is clear from global phosphoproteomes that the response is not linear, and likely relatively complex. MARK/RAF kinases now offer a strong starting point to mechanistically dissect the response pathway, including its receptor. It is encouraging that ABP1 is deeply conserved among land plants and algae (Supplementary Figure 6). While no clear ortholog is present in Marchantia^74^, ABP1 is member of the large Cupin family, and other members of this family in Arabidopsis also appear to function as auxin receptors ^75 co-submitted manuscript^. This raises the interesting possibility that the broader Cupin family, represented in all domains of life^76^, may act as auxin receptors for fast responses, including those mediated by MARK/RAF.

One striking aspect of the phosphorylation response we have discovered, is that it clearly predates the origin of the nuclear auxin response pathway^11^. Thus, well before the innovations appeared that led to auxin-dependent gene regulation, algal cells possessed a system to rapidly respond to auxin. The nuclear auxin response did not evolve to replace this system, as the rapid response system has been retained in land plants. Thus, the rapid system likely regulates responses that the nuclear system cannot, and vice versa. This could in part reflect the fundamental difference in auxin controlling cellular physiology and cell identity and fate, which happen at very different timescales. The description of this response and its deep origin, and the identification of the first component, now opens avenues to genetically and biochemically characterize these pathways in the future. This will likely deepen our understanding on the origins of auxin signaling and help reveal the ancestral role of auxin within the green lineage.

## ACKNOWLEDGEMENTS

We are grateful to Asuka Shitaku and Eri Koide for generating and sharing the Marchantia MARK/PRAF-mCitrine line, to Peng-Cheng Wang for sharing the Arabidopsis mark/raf mutants, and to our team members for discussions and helpful advice. This work was supported by funding from the Netherlands Organization for Scientific Research (NWO): VICI grant 865.14.001 and ENW-KLEIN OCENW.KLEIN.027 grants to D.W. and VENI grant VI.VENI.212.003 to A.K.; the European Resaerch Council AdG DIRNDL (contract number 833867) to D.W., CoG CATCH to J.S., StG CELLONGATE (contract 803048) to M.F. and AdG ETAP (contract 742985) to J.F.; MEXT KAKENHI Grant number JP19H05675 to T.K.; JSPS KAKENHI Grant Number JP20H03275 to R.N., Takeda Science Foundation to R.N.; the VLAG graduate school to M.v.G. and the Austrian Science Fund (FWF, P29988) to J.F..

## AUTHOR CONTRIBUTIONS

Conceptualization: A.K., M.R., D.W.; Methodology: A.K., M.R., P.C.C., S.M., S.M.D., J.S.; Formal analysis: M.R., A.K., P.C.C., S.M., S.M.D., A.M., M.F., J.S..; Investigation: A.K., M.R., P.C.C., S.M., S.M.D., A.M.; Resources: R.N., T.K.; Writing – Original Draft: A.K., D.W.; Writing – Review & Editing: all authors; Visualization: A.K., M.R., P.C.C., S.M., S.M.D.; Supervision: M.F., J.F., J.S., D.W.; Funding Acquisition: A.K., M.F., J.F., J.S., D.W..

## MATERIALS AND METHODS

### Plant material and culture conditions

All plants were cultured under 90-100 μmol photons m^-2^ s^-1^ white light with a 16 h light / 8 dark cycle at 22 °C and 75% humidity. *Arabidopsis thaliana* wild type Columbia-0 (Col-0) and all Arabidopsis mutants and transgenics were cultured on half strength Murashige and Skoog (MS) basal medium ^77^ at pH 5.7 supplemented with 0.8 % agar. All Arabidopsis mutants use were previously published: *tmk1-1* (SALK_016360) ^78^, *abp1-td1* ^79^, *afb1-3* ^80^, *mark/raf*^*null*^ (published as OK^130null^) ^58^ and *mark/raf*^*weak*^ (published as OK^130weak^) ^58^.

*Marchantia polymorpha* wild type strain Takaragaike-1 (Tak-1) and all Marchantia mutants and transgenics were cultured on half strength Gamborg’s B5 medium (B5 medium, ^81^) pH 5.7 supplemented with 1% agar. The Marchantia *mark/praf*^*ko*^ mutant was previously published as *Mppraf*^*ko* 59^.

*Klebsormidium nitens* (NIES-2285) and *Physcomitrium patens* (Gransden strain) was cultured on BCD medium ^82^ supplemented with 1 % agar under the same condition as M. polymorpha. *Penium margaritaceum* was cultured in liquid Woods Hole medium ^83^ at pH 7.2 under gentle agitation (60RPM) at 20 °C with a 16 h light / 8 dark cycle, 30 – 50 μmol photons m^-2^ s^-1^ light in 50 ml Erlenmeyer flasks.

### Phosphoproteomics

Treatment for phosphoproteomics was carried out as described in (Roosjen, Kuhn et al., accompanying manuscript) with the following adjustments: *Klebsormidium nitens, Physcomitrium patens* and *Marchantia polymorpha* were grown for 10 days on plates as described above, then treated with 100 nM IAA or DMSO in the respective growth medium for 2 minutes, harvested and frozen in liquid nitrogen. *Penium margaritaceum* was grown for 15 days as described above. Cells were collected by centrifugation at 1620 g for 2 min and washed 3 times with 10 ml of WHM to remove any residual extracellular polysaccharides from the cell surface. The pellet was resuspended in 10 ml of media and cells were treated with 100 nM IAA or DMSO for 2 min, harvested by centrifugation at 1620 g for 2 min and frozen in liquid nitrogen. Sample preparation and data analysis was carried out as described in (Roosjen, Kuhn et al., accompanying manuscript) with the following adjustments: for *Marchantia polymorpha* the UP000244005 proteome was used, for *Physcomitrium patens* the UP000006727 proteome was used, for *Klebsormidium nitens* the UP000054558 proteome was used and for *Penium margaritaceum* the proteome from a whole genome assembly was used ^48^. The mass spectrometry proteomics data, protein lists and intensity values of all samples have been deposited to the ProteomeXchange Consortium via the PRIDE ^84^ partner repository with the dataset identifier XXX. All phosphoproteomics data has been compiled in the AuxPhos web-app (https://weijerslab.shinyapps.io/AuxPhos; Roosjen, Kuhn et al., accompanying manuscript).

### Orthogroup construction

Identification of orthogroups i.e., common orthologous sequences between multiple species were estimated using Orthofinder ^85^. Proteomes used for this analysis include: *Arabidopsis thaliana* (Araport11), *Marchantia polymorpha* (v6.1), *Physcomitrium patens* (v3.3), *Klebsormidium nitens* (v1.1) and *Penium margaritaceum* (v1).

### Cytoplasmic streaming

Cytoplasmic streaming was recorded using a Leica SP5 or SP8 confocal microscope equipped with HyD detectors using Apo λ 63×/1.10 water immersion objective plus 6x digital zoom in an 256×256 pixel format. Cytoplasmic streaming was recorded and analyzed for Arabidopsis epidermal cells of the root elongation zone and Marchantia rhizoid cells using the following method: Seven day old Arabidopsis plate-grown seedlings were taken into the microscopy room and mitochondria were stained by transferring the seedling into a petri dish with liquid ½ MS medium containing 1 μM Rhodamine 123 for 5 minutes. Subsequently, seedlings were washed with liquid ½ MS without Rhodamine 123. Seedlings were then transferred to microcopy slides in a drop of liquid ½ MS containing 100 nM IAA or DMSO, covered by a coverslip and left on the microscope stage to adapt to the environment for 30 minutes. Cytoplasmic streaming was recorded in at least 5 epidermal cells of the root elongation zone per root at a frame rate of 5.3 frames per second for 30 seconds (159 frames).

Prior to the experiment, Marchantia thallus was grown from gemmae for two days in liquid B5 medium in a petridish. After two days of cultivation Rhodamine 123 was added to a final concentration of 1 μM and Triton-X-100 was added to a final concentration of 0.01%. Marchantia samples were stained for 30 minutes and then washed three time with liquid B5 medium containing 0.01% Triton-X-100 without Rhodamine 123. Samples were then transferred to microscopy slides and cytoplasmic streaming in rhizoid cells was recorded as described for Arabidopsis.

### Data analysis for cytoplasmic streaming

Data analysis was performed in MatLab (version: 2021b). First, static background signal was removed from the raw fluorescence images using a moving window median filter (averaging window = 25 frames) and motile objects smoothed with a 2-pixel Gaussian blur filter. Moving objects were tracked using an established particle tracking algorithm ^86^, keeping only those trajectories whose length exceeds 3 seconds. For each cell, from the individual trajectories of the remaining moving objects, typically between 30 to 60 per time series, an ensemble-averaged mean-squared displacement was computed:

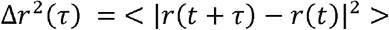

Per cell, these mean-squared displacements were fitted to the anomalous diffusion model (ADM) ^87,88^, a generalization of Einstein’s diffusion model to describe complex non-Fickian motion of organelles in the visco-elastic liquid of the cellular cytosol, which is composed of an unknown mixture of passive (Brownian) and active (streaming) transport in a crowded and heterogeneous medium:

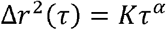

where *π* is the correlation time. In the ADM, the generalized diffusion power law exponent *α*provides information on the average nature of the transport processes: *α*<1 is indicative of sub-diffusive motion, characteristic of Brownian motion in a visco-elastic liquid, *α* = 1 indicates pure Brownian motion in a viscous liquid and *α* > 1, known as super-diffusion, indicates transport with an active, e.g., motor-protein driven, component. Intermediate values of the power law exponent *α*provide insight into the relative balance of these different processes on the organellar motion. The transport rate constant *K* (in units mm/s^a^) informs about the average transport rate: the larger the value of *K* the faster the organellar transport in the cells. Our analysis yields one average value for *α*and *K* per cell; the significance of the differences between control and treatment was assessed with a two-sided Wilcoxon signed rank test.

### Membrane potential measurement using DISBAC_2_(3)

Membrane potential was measured using the DISBAC_2_(3) probe as previously described for Arabidopsis^17^. DISBAC2(3) (2 μM) was added to buffered ½ MS liquid medium with 1% (w/v) sucrose containing either 0 or 100 nM IAA. Five-day-old Arabidopsis seedlings were transferred to a sealable single-layer PDMS silicone chip^17^. The PDMS silicone chip containing the seedlings was then placed on a vertical spinning disk microscope for a 20-min recovery. During the recovery process, the seedlings were treated with control medium at a flow rate of 3 μl/min. Seedlings were imaged every 30 seconds with a x20/0.8 objective. DISBAC2(3) was excited with a 515-nm laser, and the emission was filtered with a 535/30-nm bandpass filter. DISBAC2(3) fluorescence was measured at the border between epidermis and cortical cells of the transition zone by selecting 5-6 or 3-4 cells for Col-0 and *Atmark/raf*^*null*^, respectively.

Membrane potential of Marchantia and Klebsormidium was measured using the same probe with the following modifications to the protocol: Marchantia gemmae were removed from gemmae cups and placed liquid B5 with 0.01%Triton-X-100 supplemented with 15 μM DISBAC_2_(3), vacuum infiltrated for 5 minutes and transferred to a cover slip followed by incubation for 30 minutes before imaging. Imaging was performed on an inverted Leica SP8 confocal microscope using the same setting as for Arabidopsis. Klebsormidium was grown for 10 days as described above. A small amount of Klebsormidium was then scraped off the plate and dissolved in liquid BCD medium supplemented with 15 μM DISBAC_2_(3) followed by incubation for 30 minutes before imaging.

### Root surface pH profile

Root surface pH was measured using the ratiometric Fluorescein-5-(and-6)-Sulfonic Acid, Trisodium Salt (FS) (Invitrogen™ F1130)^89^. Five-day-old Arabidopsis seedlings were transferred to unbuffered ½ MS medium containing 50 μM FS dye and either 0 or 100 nM IAA. Seedlings were allowed to recover on a vertical spinning disk microscope for 20 minutes after transfer to the microscope chamber. Imaging was performed using a vertical stage Zeiss Axio Observer 7 microscope coupled to a Yokogawa CSU-W1-T2 spinning disk unit with 50 μm pinholes, equipped with a VS-HOM1000 excitation light homogenizer (Visitron Systems). Images were acquired using VisiView software (Visitron Systems, v.4.4.0.14). We used a Zeiss Plan-Apochromat ×10/0.45 objective. FS was excited by 405 and 488 nm laser. The 488/405 nm fluorescence emission ratio along the root was calculated using the ATR software^89^.

### Phenotyping

Arabidopsis ***plant height*** was determined from respectively 48 individual wild type and *mark/raf*^*null*^ senescing plants, seven weeks after germination. To compare the ***leaf area*** of fully elongated leaf 6 to leaf 9 of Arabidopsis wild type, *mark/raf*^*null*^ and *mark/raf*^*weak*^, 16 plants per genotype were collected, flattened on paper and photographed using a Canon EOS 250D with EFS 18-135mm Macro Lens. Leaf area was measured in ImageJ (Version 1.52) using the Polygon selection tool.

***Rosette area*** was determined from respectively 90 individual wild type and *mark/raf*^*null*^ plants plants 28 days after germination. Plants were photographed individually using a Canon EOS 250D camera with EFS 18-135mm Macro Lens. Rosette area was then measured in ImageJ (Version 1.52) using the Polygon selection tool. To compare the ***germination efficiency*** between *mark/raf*^*null*^ mutants and wild type, seeds for each genotype were surface sterilized, stratified in a 0.1% agarose solution for two days at 4 °C and paced on half strength MS plates (0.8% Agar). Plates were grown vertically for 9 days and germinated seeds were scored at day 1, 2, 3, 4, 7 and 9. Germination percentages were calculated for each day. The experiment was repeated three times individually and data were combined for analysis.

Seedlings of Arabidopsis wild type, *mark/raf*^*null*^ and *mark/raf*^*weak*^ were germinated on half strength MS and vertically grown for 5 days. After five days, ten seedlings with representative root length for each genotype were transferred to new square petri dishes either containing 1 nM IAA, 100 nM IAA or a mock treatment representing an equal amount of solvent (DMSO). ***Root length*** was captured by photographing the plates immediately after transferring the seedlings, after 24 hour, after 48 hours and after 120 hours, using a Canon EOS 250D camera with EFS 18-135mm Macro Lens. Root length was then measured in ImageJ using the segmented line tool and growth rates calculated.

To compare the ***thallus growth*** between Marchantia *mark/praf*^*ko*^ mutants (n=44) and wild type (n=50), thalli were grown from gemmae on half strength Gamborg B5 medium. Plates were grown for 29 days and projected thallus area was captured by photographing the plates immediately after transferring the gemmae, after 2, 4, 7, 9, 11, 14, 16, 18, 22 and 29 days, using a Canon EOS 250D camera with EFS 18-135mm Macro Lens. Thallus area was then measured in ImageJ (Version 1.52) using the Polygon selection tool. For ***auxin sensitivity*** assays, Marchantia *mark/praf*^*ko*^ mutant (n=10) or wild-type (n=10) gemmae were grown on half strength Gamborg B5 medium supplemented the indicated concentration of IAA and grown for 10 days. At day 10, thallus size was captured by photographing the plates using a Canon EOS 250D with EFS 18-135mm Macro Lens. Thallus area was then measured in ImageJ (Version 1.52) using the Polygon selection tool. ***Gemma cup number*** was determined on *mark/praf*^*ko*^ mutants (n=14) and wild type (n=14) thalli after 24 days of growth on half strength Gamborg B5 medium.

### Transcriptomic analysis

*Arabidopsis thaliana* wild-type (Col-0) and mutant (*mark/raf*^*null*^) seeds were sown on half-strength MS medium covered with nylon mesh and vertically grown for 7 days. Plants were then submerged in liquid half-strength MS medium containing either 1 μM IAA or the equivalent amount of solvent (DMSO). Plates were kept horizontally for about 30 seconds and then kept vertically for 1 hour to incubate. After incubation, root tips were harvested using a scalpel and immediately frozen in liquid nitrogen.

*Marchantia polymorpha* wild-type (Tak-1) and mutant (*mark/praf*^*ko*^) gemmae were placed on B5 solid medium covered with nylon mesh (100 mm pore) and grown for 9 days. After growing, plants were submerged in liquid B5 medium and cultured for 1 day. After pre-cultivation, IAA was added to a final concentration of 1 μM or an equivalent amount of DMSO was added and plants were incubated for 1 hour. Using a scalpel, thalli were harvested from the mesh, blotted on paper towels and immediately frozen in liquid nitrogen.

After harvesting, all frozen samples were ground into fine powder using a pre-cooled mortar and pestle. Total RNA form all samples was extracted using a RNeasy Plant Mini Kit (QIAGEN). Total RNA was treated with RNase-free DNase I set (QIAGEN). RNA-seq library construction and RNA sequencing were performed by BGI Tech Solutions (Hong Kong).

### RNAseq data analysis

Up to 20 million paired-end 150 bp reads were collected for each sample. Quality assessment for raw reads was performed using FastQC (www.bioinformatics.babraham.ac.uk/projects/fastqc). For both *Arabidopsis thaliana* (Araport11; ^90^) and *Marchantia polymorpha* (v6.1; ^91^), reads were mapped onto the respective genomes using HISAT2 (v2.1.0; ^92^) with additional parameters “--trim5 10 –dta”. Alignment (SAM/BAM) files were sorted and indexed using SAMTOOLS (v1.9;^93^). FeatureCounts (v2.0.0; ^94^) was used to count the reads mapped on to each gene, with the parameters “-t ‘exon’ -g ‘gene_id’ -Q 30 --primary -p -B -C” for Arabidopsis transcipts and “t ‘gene’ -g ‘ID’ -Q 30 --primary -p -B -C” for Marchantia transcripts. DEseq2 ^95^ was used to normalize the raw counts and perform the differential expression analysis with a design matrix including the interaction term (Padj<0.05). Data processing and statistical analysis was performed using R (https://www.r-project.org/). Sequenced raw reads were deposited in NCBI Sequence Read Archive (SRA) under the project accession number PRJNA881051.

### Generation of transgenics

Primers used in this study can be found in Supplementary Table 1. Arabidopsis MARK reporter lines for MARK1/RAF24 and MARK5/RAF20 under their endogenous promoter were generated by amplifying the genomic fragment including the 3.5 kb region upstream of the start codon using the appropriate primers for each gene. Fragments were cloned into a pGIIK LIC-YFP (pPLV17) vector ^96^ using the HiFi cloning kit (ThermoFisher).

For the Marchantia MARK/PRAF reporter line, a DNA fragment for an Arabidopsis-codon-optimized mCitrine coding sequence (CDS) was synthesized (IDT) and used to amplify a GGSC2 linker-containing fragment by PCR with a primer set, pUGW2_Aor_GGS2_mCit_IF_F and pUGW2_Aor_mCit_IF_R, which was then cloned into the Aor51HI site in pUGW2 35S ^97^ using the In-Fusion cloning kit (TaKaRa Bio). The 2.5-kb HindIII-SacI fragment in the resulting plasmid, including the Gateway cassette followed by the GGSC2 linker-attached mCitrine CDS, was ligated with the HindIII- and SacI-digested pMpGWBx00 ^97^ to generate pMpGWBx47. The MpMARK/PRAF genomic sequence covering its promoter and CDS (without stop codon) in pENTR/D-TOPO_MpMARK/PRAF ^59^ was transferred to pMpGWB347 to generate pMpGWB347-MpMARK/PRAF. Agrobacterium GV2260 containing pMpGWB347-MpMARK/PRAF was used to transform Mpmark/praf^ko^ plants (Koide et al. 2020) by the thallus transformation method ^98^.

### Imaging of transgenic lines plants for MARK-localization analysis

Marchantia gemmae expressing MARK/PRAF-mCitrine under endogenous promoter and 7 day-old Arabidopsis roots expressing MARK1/RAF24-YFP or MARK5/RAF20-YFP under their respective endogenous promoter were imaged using a Leica SP5 or SP8 confocal microscope equipped with an Argon laser (SP5) or a white light laser (SP8). Both, mCitrine and YFP were excited at 514 nm, and emission was collected between 525-575 nm. Images were analyzed using ImageJ (Version 1.52).

## FIGURE LEGENDS

**Supplementary figure 1. Cytoplasmic streaming relies on the actin cytoskeleton** (A,B) Quantification of the diffusive component (α) of cytoplasmic streaming in wild-type and *mark/raf* mutant Arabidopsis roots (**A**) and wild-type and Atmark/praf mutant Marchantia thallus (**B**) with and without auxin treatment. (C,D) Diffusive Exponent (α; C) and Diffusion Rate (K; D) of cytoplasmic streaming in wild-type Arabidopsis roots treated with mock medium or Lantrunculin B. Boxplots are shown along individual measurements, number of observations (n) is indicated, and significance (Student’s t-test) is shown.

**Supplementary figure 2 Phenotypic analysis of *mark* mutants in Marchantia and Arabidopsis**

(**A**) Projected thallus area in wild-type and *Mpmark/praf*^*ko*^ mutant Marchantia thallus, followed over 29 days. **(B)** Number of gemma cup on wild-type and *Mpmark/praf*^*ko*^ mutant Marchantia thallus. **(C)**. Root length in wild-type and *mark/raf*^*null*^ mutant Arabidopsis seedlings, followed over 9 days. **(D**,**E)** Rosette area (**D**) and height (**E**) in wild-type and *mark/raf*^*null*^ mutant Arabidopsis plants. **(F)** Germination rate of wild-type and *mark/raf*^*null*^ mutant Arabidopsis seeds, followed over 9 days. (**G**,**H**) Examples of images used for quantification in panel **A** and **C**, respectively.

**Supplementary figure 3: Analysis of *mark* mutant phosphoproteomes**.

**(A)** Overlap of MARK targets in Arabidopsis and Marchantia, based on differential phosphorylation in *Atmark/raf* and *Mpmark/praf* phosphoproteomes under mock conditions, compared to wild-types. **(B)** Venn diagram showing overlap between phosphosites differentially regulated (≤0.05) in mannitol-treated Arabidopsis plants (Lin et.al. 2020), 100nM IAA treated Col-0 and 100nM treated *mark* null mutant. (**C**) Gene identifiers of Arabidopsis MARK/Raf kinases.

**Supplementary Figure 4: Dynamics of membrane potential in wild-type and *mark/raf* mutant Arabidopsis roots**

**(A)** Arabidopsis wildtype (Col-0) and *mark/raf*^*null*^ mutant root surface pH visualized using the ratiometric pH-sensitive FS dye treated with mock or 100 nM IAA; scale bar = 50 μm. **(B)** Quantification of the F488/405 nm fluorescence emission ratio along the root surface of wildtype (Col-0) and *mark/raf*^*null*^. Higher ratio corresponds to alkaline pH. Control and 100 nM IAA-treated roots are shown. The graphs show the averages 12 and 11 roots for wildtype (Col-0) and *mark/raf*^*null*^, respectively for both mock and IAA conditions. Shaded areas represent standard deviations. (**C**) Dynamics of membrane potential after treatment with 100 nM IAA (arrow) in *Atmark/raf*^*null*^ (n=10), Col-0 (n=6) and *afb1-3* (n=6) roots. Membrane potential was visualized by the relative change of the DISBAC2(3) fluorescence over time in a microfluidic chip. Average values are shown, shaded areas represent standard deviations.

**Supplementary figure 6 Phylogenetic analysis of ABP1**.

**(A)** Phylogenetic tree of the ABP1 gene family with green algae and land plant homologs. Branches that are well-supported (bootstrap >75) are marked with dots. Orthologs from each phylum are represented with a different color. **(B)** Deep conservation of key amino acids in the ABP1 auxin binding pocket, as well as the Zinc binding site. Light blue to dark blue color gradient represents low to high conservation, respectively. Numbering on the top is based on maize ABP1 protein ^99^. The complete tree can be found at interactive Tree of Life (iTOL): https://itol.embl.de/shared/dolfweijers.

