## Supplementary figures and images for "A RAF-like kinase mediates a deeply conserved, ultra-rapid auxin response"

### Figure S1

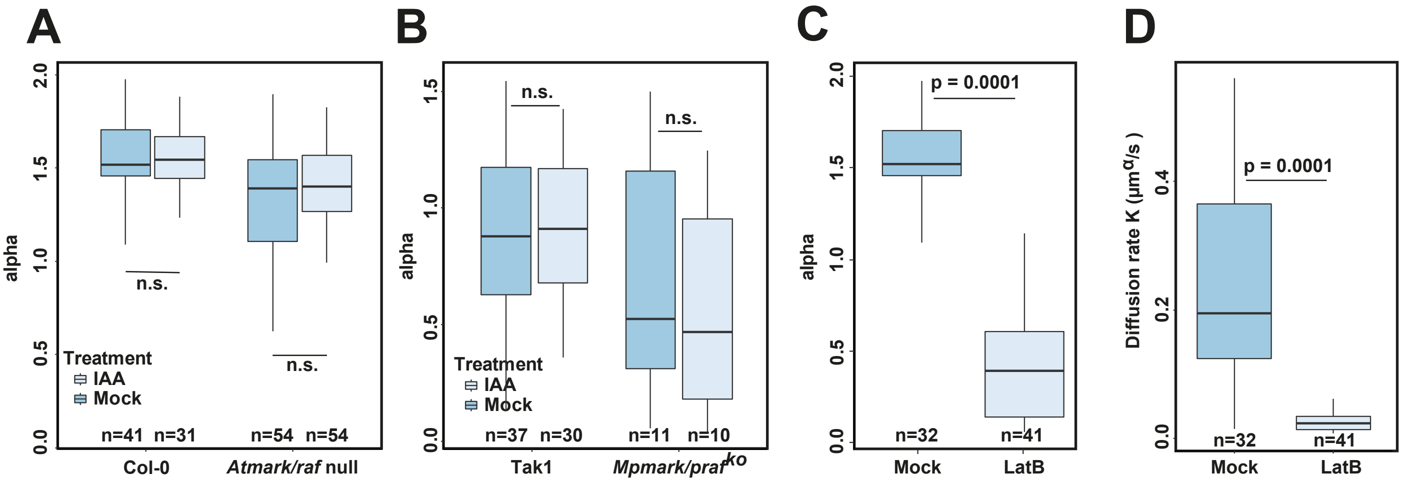

### Figure S2

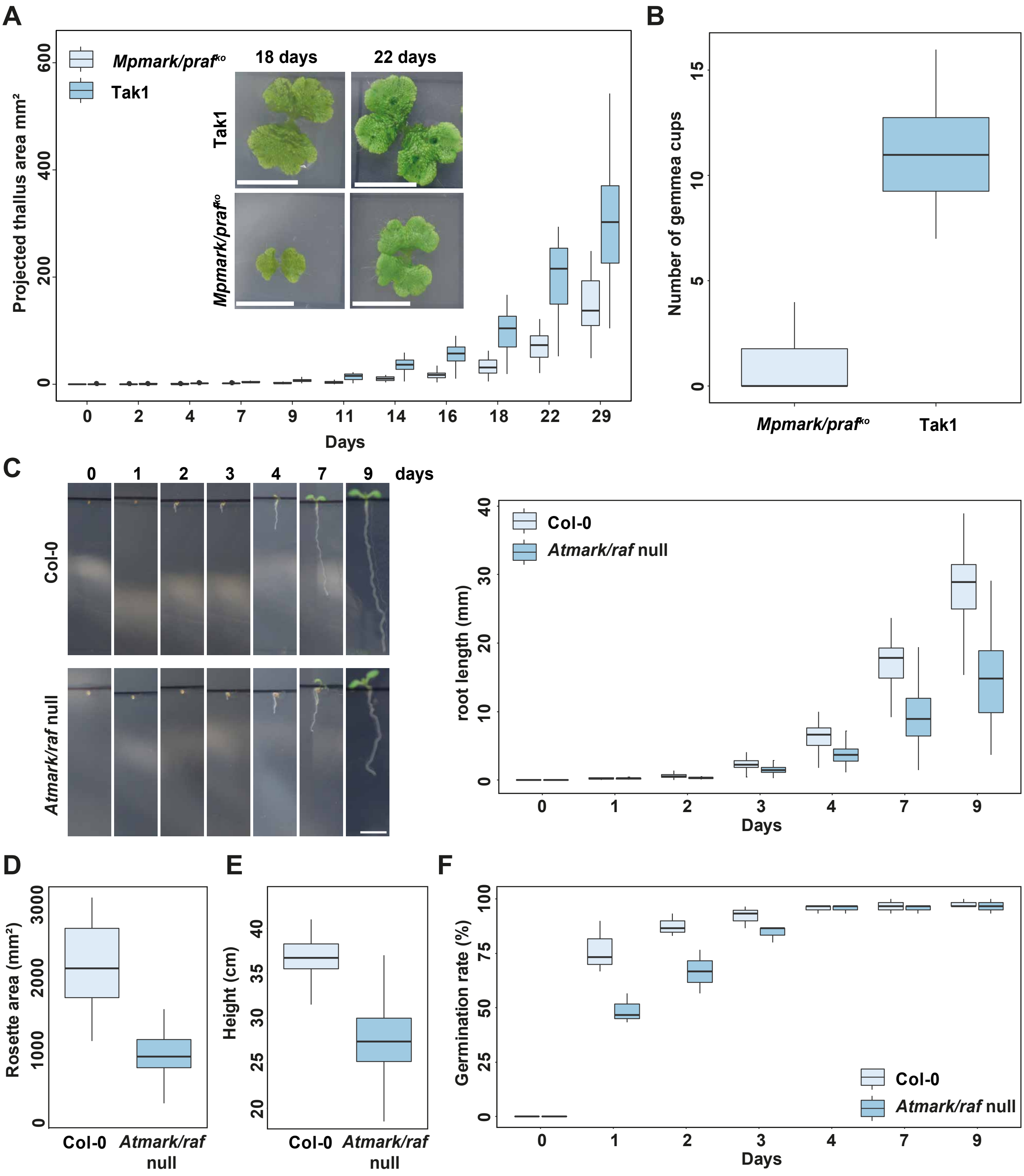

### Figure S4

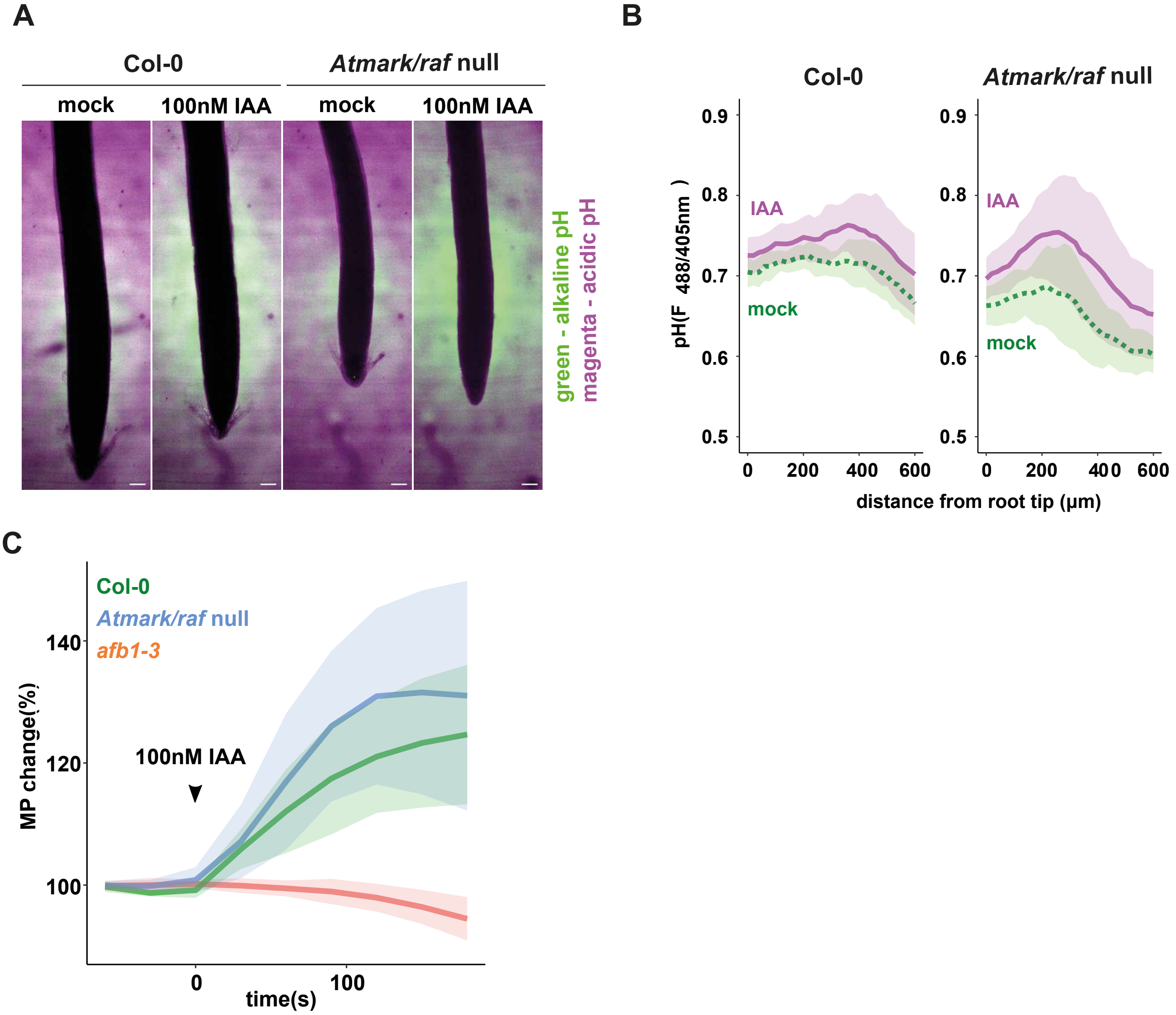

### Figure S5

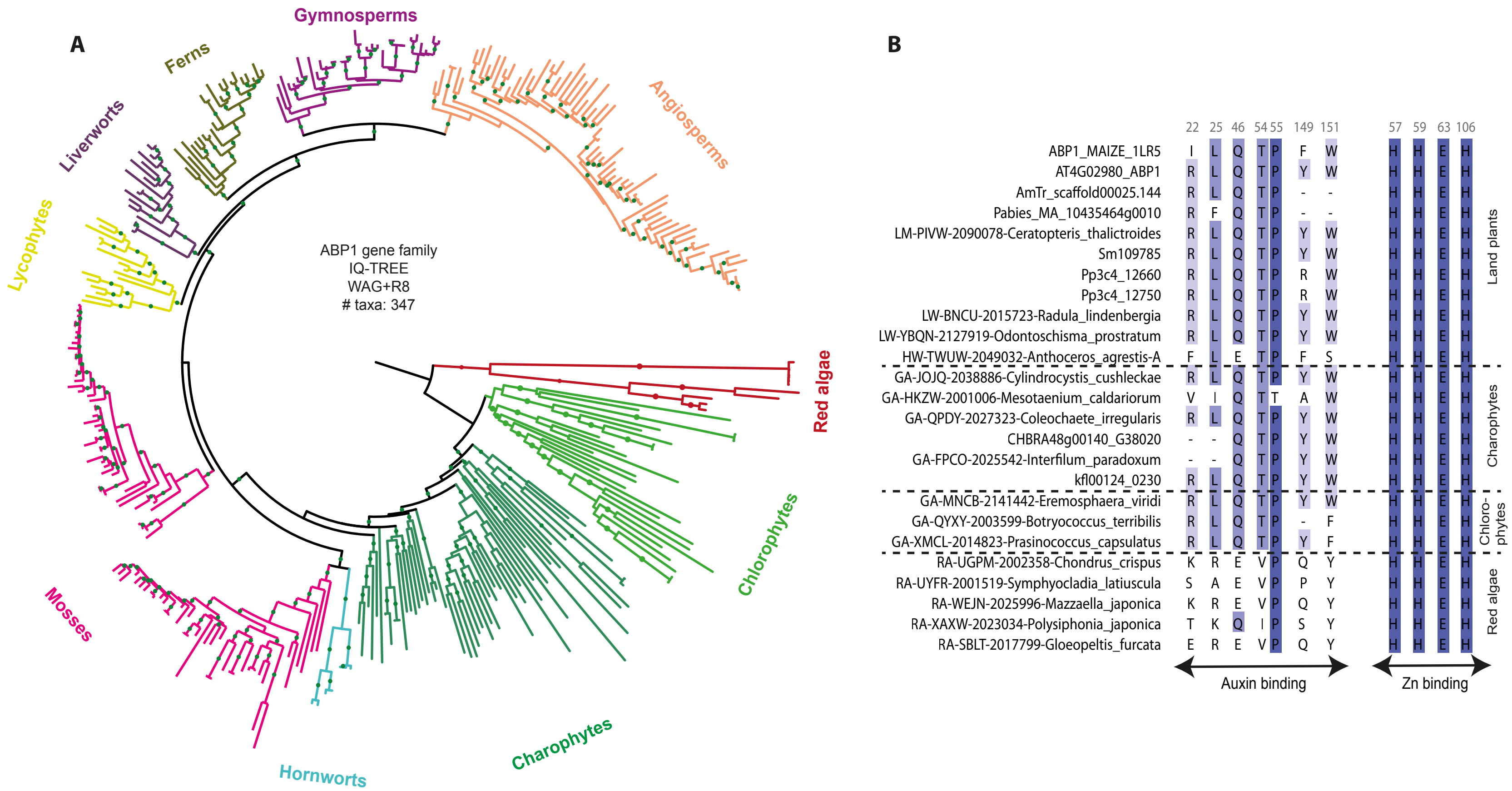
