## Supplementary material for "A RAF-like kinase mediates a deeply conserved, ultra-rapid auxin response": Figure S3

A

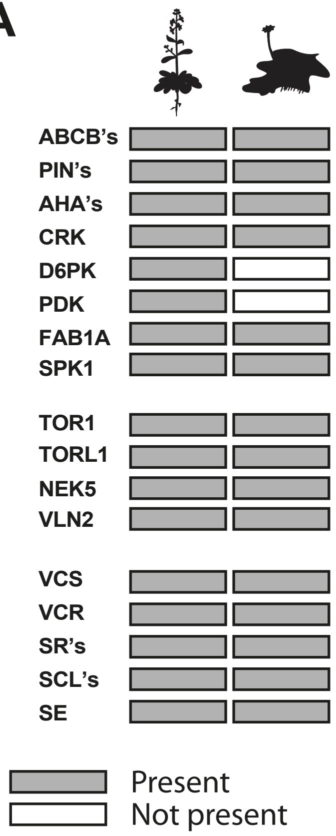


B

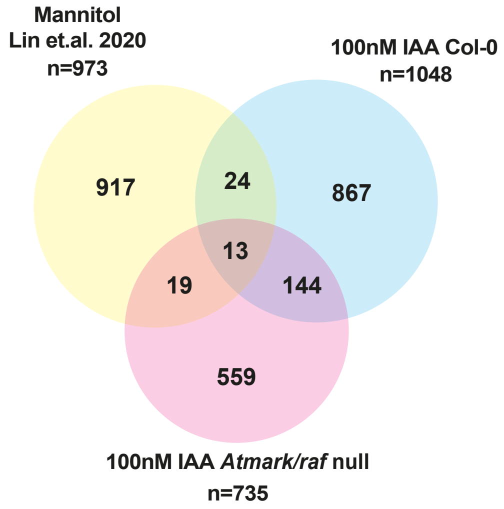


C

| ATG Code | MARK Code | RAF Code |
|---|---|---|
| AT2G35050 | MARK1 | RAF24 |
| AT5G57610 | MARK2 | RAF35 |
| AT1G04700 | MARK3 | RAF16 |
| AT1G16270 | MARK4 | RAF18 |
| AT1G79570 | MARK5 | RAF20 |
| AT3G46920 | MARK6 | RAF42 |
| AT3G24715 | MARK7 | RAF40/HCR1 |
